# Differential mRNA localization of *karyopherin-β*2 homologs in *C. elegans* and humans

**DOI:** 10.64898/2026.05.19.726232

**Authors:** Ambika Basu, Nora Tayefeh, Lindsay P Winkenbach, Erin Osborne Nishimura

## Abstract

In *Caenorhabditis elegans* embryos, the nuclear transport receptor IMB-2 (Importin Beta Family-2, a karyopherin *β*2) preferentially localizes to the nuclear envelope along with its encoding mRNA. This suggests that *imb-2* mRNA is locally translated at the nuclear envelope. To test whether *imb-2*’s two putative human orthologs, Transportin 1 (TNPO1) and Transportin 2 (TNPO2), exhibited similar mRNA localization, we performed smiFISH and microscopy in U2OS, HeLa, and human pluripotent stem cells. Neither human *TNPO1* nor *TNPO2* mRNA localized to the nuclear envelope in any tested human cell type. However, the human TNPO1 protein and the *C. elegans* IMB-2 protein both localized to the nucleus and its periphery. This suggests that despite their shared functional roles these karyopherins have different routes to expression.

**Article summary:** Gene expression is a multistep process. Genes are transcribed into mRNA in nuclei and translated into proteins in the cytoplasm. *C. elegans imb-2* localizes preferentially at the nuclear envelope for reasons that are unclear. We asked whether human *imb-2* homologs also localize at the nuclear envelope. Two *imb-2* orthologs in humans, Transportin-1 and 2, showed no preference for the nuclear envelope. However, once translated, they all shared the same nuclear localization and all function to move proteins in and out. This work will appeal to geneticists, cell biologists, and molecular biologists.

## Introduction

Transcription canonically occurs in the cytoplasm or, for secreted proteins, on the rough endoplasmic reticulum. However, translation can also occur at diverse subcellular locales (Lécuyer *et al*. 2009; Lasko 2012; Mofatteh 2020; Das *et al*. 2021; Lashkevich and Dmitriev 2021). Early evidence of spatially specific translation, termed “local translation”, emerged out of observations of mRNA localization during the oocyte-to-embryo transition, such as in sea squirt embryos (Jeffery *et al*. 1983), Xenopus oocytes (Rebagliati *et al*. 1985), and in the Drosophila embryonic syncytium (Anderson *et al*. 1985; Berleth *et al*. 1988). mRNA localization and local translation were initially hypothesized to be specialized behaviors of oocytes, embryos, and neurons, in which local translation was important for the oocyte-to-embryo transition or required for transport down long neuronal projections (Johnston 1995; Lécuyer *et al*. 2009). Originally, mRNA localization and local translation were not expected to be a general feature of cell biology.

Over the decades, numerous examples of mRNA localization and local translation have emerged in a wide array of non-neuronal and non-embryonic systems (Long *et al*. 1997; Moor *et al*. 2017; Chouaib *et al*. 2020; Lashkevich and Dmitriev 2021). For example, bacteria accumulate mRNAs and sRNAs (small regulatory RNAs) at diverse sites (Fei and Sharma 2019; Shang *et al*. 2024). In metazoans, mRNA localization has been reported at a wide array of subcellular locales, such as at the endoplasmic reticulum (ER) (Akopian *et al*. 2013; Ast *et al*. 2013), in apical or basal compartments of polarized cells (Moor *et al*. 2017; Novoselsky *et al*. 2026), at the mitochondrial membrane (Fazal *et al*. 2019; Müntjes *et al*. 2021), near pericentriolar material (Sepulveda *et al*. 2018; Kim *et al*. 2019; Zein-Sabatto *et al*. 2024), at the plasma membrane (Wu *et al*. 2021), at cell-to-cell junctions (Chin *et al*. 2026), in TIS granules (Ma and Mayr 2018), and within a number of different biomolecular condensates (Banani *et al*. 2017; Seydoux 2018; Trcek and Lehmann 2019; Parker *et al*. 2022). These discoveries have arisen from advances in RNA methods such as proximity labeling that captures subcellular transcriptomes (Fazal *et al*. 2019; Padrón *et al*. 2019; Lo *et al*. 2022), imaging methods (smFISH and fluorescent aptamers) (Raj *et al*. 2010; Shaffer *et al*. 2013; Tsanov *et al*. 2016), RNA immunoprecipitation (Wheeler *et al*. 2018), and subcellular fractionation (Khong *et al*. 2017). Indeed, mRNA localization and local translation appear more widespread than initially envisioned.

How conserved is local translation across metazoans? Some sub-cellular locales seem particularly prone to local translation, such as cell junctions and synapses (Li *et al*. 2021; Tocchini *et al*. 2021; Tu *et al*. 2024; Chin *et al*. 2026). However, it is not clear whether local translation is conserved among direct gene orthologs. That is, we do not know whether a protein’s local translation in one organism predicts its local translation in another. Therefore, identifying conserved examples of local translation across species will be important to better understand translation evolution and the biology of the underlying protein. By focusing research on the most conserved examples of local translation, we can ensure that model-organism research yields the greatest potential positive impact on human health and leverages an array of organism-specific technologies.

Our group reported that the *C. elegans* IMB-2 protein is translated from a localized mRNA transcript concentrating at the nuclear envelope (Parker *et al*. 2020). Despite its name, *C. elegans* IMB-2 shares a closer sequence and domain similarity to human Transportin 1 (also known as karyopherin-*β*2, TNPO1, and TRN1) and Transportin 2 (also known as karyopherin-*β*2*β*, TNPO2, and TRN2) than to human Importin-*β* (Quan *et al*. 2008; O’Reilly *et al*. 2011; Yang *et al*. 2023). The transportin/karyopherin-*β*2 subfamily is comprised of karyopherins that usher protein cargoes from the cytoplasm to the nucleus, directed by Nuclear Localization Sequences (NLSs) in their protein passengers (Mboukou *et al*. 2021). Unlike the canonical Importin-*β* pathway, Transportins do not require heterodimerization with Importin-*α* to function and they recognize a “non-classical” NLS. Transportin cargo import is mediated by interactions with nuclear pore complex subunits, RanGTP, and sometimes with other transport proteins (Mboukou *et al*. 2021). In *C. elegans*, IMB-2 is required to transport key cargos such as DAF-16 to mediate stress response and aging (Senchuk *et al*. 2018), and NSY-7, a transcription factor required for stochastic olfactory neuron development (Alqadah *et al*. 2019).

Because karyopherins like IMB-2 are conserved across metazoans, we set out to test whether IMB-2’s human orthologs share its characteristic mRNA localization and local translation. The *C. elegans* IMB-2 protein has 52% amino acid identity with TNPO1 and 51% with TNPO2 (Fig. 1a, Figure S1). They also share key domains such as their N-terminal importin-*β* binding domain (IBB) and a series of tandem HEAT repeats (Fig. 1b).

**Figure 1.**
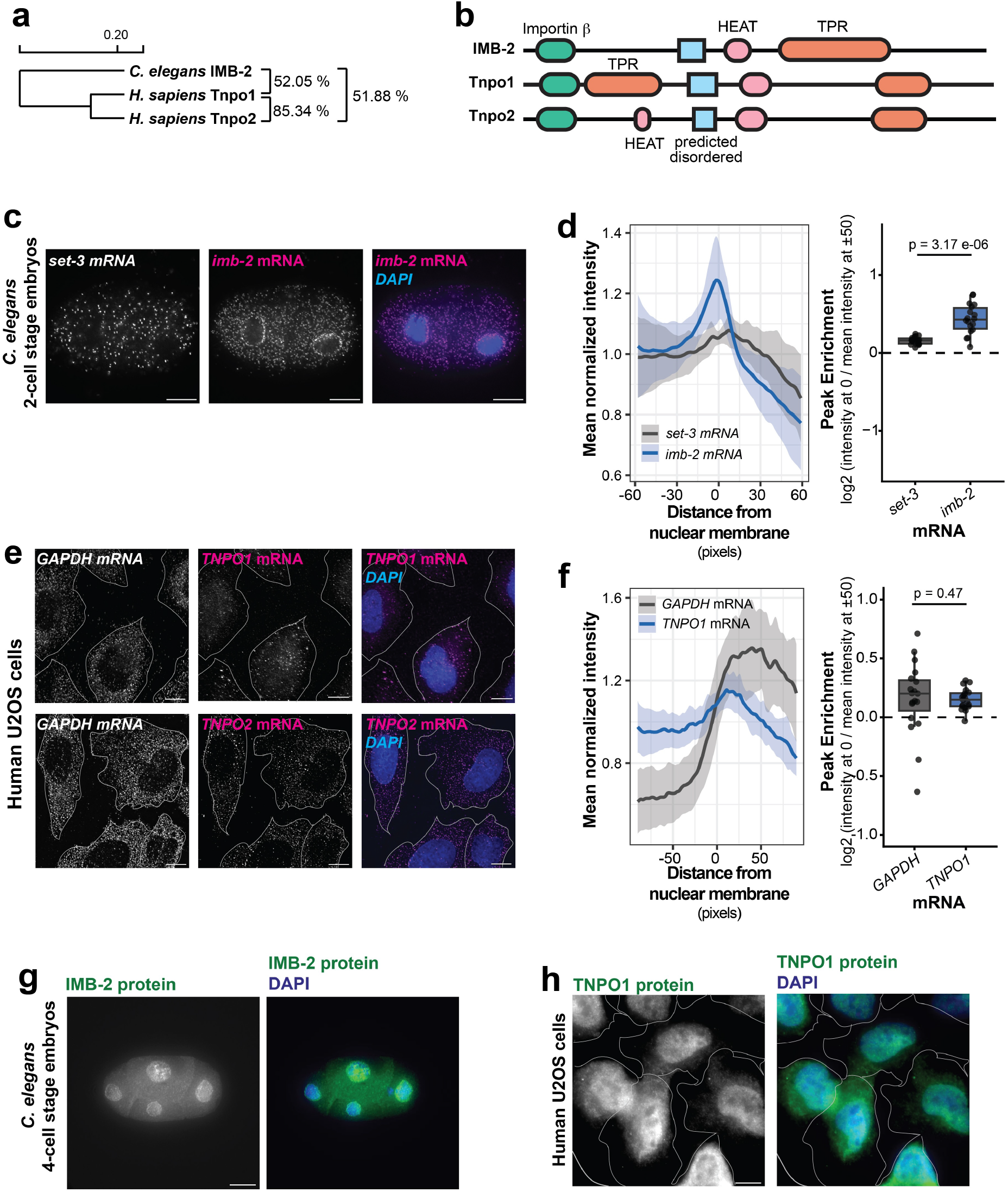
*C. elegans imb-2* and human *TNPO1* and *TNPO2* mRNA and protein expression patterns. **a**. Conservation between three transportin proteins: human TNPO1, human TNPO2, and *C. elegans* IMB-2. Sequence identity is plotted as a dendrogram. Branch lengths are proportional to changes. **b**. Protein domain predictions for IMB-2, TNPO1, and TNPO2 from InterPro. Importin beta = Importin beta N-terminal domain; HEAT = HEAT repeat; TPR = Tetratricopeptide repeats. **c**. Representative micrographs showing *C. elegans* homogenously distributed *set-3* mRNA as imaged by smFISH (white), *imb-2* mRNA as imaged by smFISH (magenta) and DAPI (blue). Bars 10 *µ*m. **d** Quantification from **b** were calculated by integrating the total signal intensity of smFISH fluorescence across a 3-D box drawn to span the nuclear membrane at x = 0 (negative x values are within the nucleus), then normalized with respect to individual-embryo-means, then averaged across samples with respect to the x-axis. We calculated a metric of fluorescence-intensity enrichment at the nuclear membrane by dividing the mean normalized intensity at x = 0 by the average between x = -50 and +50. n = 18; reps = 4. **e**. Representative images of *TNPO1* mRNA (magenta, smiFISH, top), *TNPO1* mRNA (magenta, smiFISH, bottom) and respective nuclei (blue, DAPI) with *GAPDH* mRNA (white, smiFISH) included as a homogenous control in U2OS cells. Bars 10 *µ*m. f. Quantification was performed for the samples in **e** as performed in **d**. n = 19; reps = 3. **g**. Representative micrographs showing *C. elegans* IMB-2 as imaged by GFP-tagged IMB-2 protein (green) (strain WON102 containing imb-2::GFP), and nuclei (blue, DAPI) in 4-cell stage nematode embryos. Bars 10 *µ*m. **h**.Representative images of *H. sapiens* TNPO1 protein (green, ICC) and nuclei (blue, DAPI) in U2OS cells. Bars 10 *µ*m.

Overall, mRNA localization at the nuclear envelope has been reported rarely, with fewer published examples than other subcellular locales. Key known examples are *C. elegans imb-2* (Dylan M Parker et al. 2020), human ASPM (during interphase) (Chouaib *et al*. 2020), SPEN (Chouaib *et al*. 2020), and nuclear pore complex components identified in yeast (Lautier *et al*. 2021). Therefore, *C. elegans* IMB-2 represents a valuable entry point into this emerging field.

## Materials and methods

### Cell Culture

U2OS and HeLa cells were maintained in DMEM (Dulbecco’s Modified Eagle Medium, High glucose with L-glutamine and sodium pyruvate, ThermoFisher, 11995040) supplemented with 10% fetal bovine serum and 1% penicillin in a humidified 37° incubator with 5% CO2. Nearly 20-80% cell density was maintained. Human Pluripotent Stem Cells (hPSCs) were maintained in ExCellerate iPSC Expansion Medium (R&D Systems) with 100 units/mL HyClone Antibiotic Antimycotic (Cytiva) on plates coated with 0.1 mg/mL Cultrex UltiMatrix RGF BME (R&D Systems). The cells were cultured at 37° in 5% CO2 with daily medium changes. hPSCs were passaged at 70% confluency (roughly every 4 days) after dissociation into a single cell suspension using ACCUMAX (STEM-CELL Technologies).

### GFP tagged IMB-2 protein *C. elegans* strain generation

GFP tagged IMB-2 protein *C. elegans* strain generation The GFP tagged imb-2 strain WON102 was generated by InVivo Biosystems. It was made using MosSCI transgenesis single copy insertion of *imb-2p::imb-2::GFP::imb-2 3’utr* in cxTi10882 MosSCI locus on chromosome IV.

### *C. elegans* husbandry

*C. elegans* strains were maintained and grown on nematode growth medium (NGM, 3 g/L NaCL; 17 g/L agar; 2.5 g/L peptone; 5 mg/L cholesterol; 1 mM MgSO4; 2.7 g/L KH2PO4; 0.89 g/L K2HPO4) at 20°, using standard worm husbandry practices, feeding them OP50 *E*.*coli* (Brenner 1974). To visualize GFP fusions of the IMB-2 protein, we used WON102 onsSi870 [*pNU2550 (imb-2p::imb-2::GFP::imb-2u, unc-119(+))*] IV; *unc-119(ed3)* III (Fig 1B, C). To visualize discrete cells in larval stages, we used strain ML1651 (*mcls46 [dlg-1::RFP + unc-119(+)]*) (Diogon *et al*. 2007) in which RFP marks epithelial junctions. Strains used in this study are included in 1.

### smFISH in *C. elegans* embryos

To visualize RNAs at single-molecule resolution, smFISH was used according to the protocol described in Parker et al (Parker *et al*. 2021). The Stellaris RNA FISH probe designer was used to design custom smFISH probesets. Gravid worms were bleached to harvest embryos. These were fixed and permeabilized by suspending in 1 ml -20° methanol for 4 min, followed by freeze-cracking in liquid nitrogen for 1 min, and incubation in -20° acetone for 5 min. After fixation, embryos were incubated in Stellaris Wash Buffer A (Biosearch Technologies, SMF-WA1-60) at room temperature for 5 min, then hybridized in Sterallis Hybridization Buffer (Biosearch Technologies, SMF-HB1-10) and 10% formamide with 50 pmol of each primer set according to the procedure described in (Parker *et al*. 2021). The hybridization reaction was incubated for 16-48 hr at 37° mixing at 400 rpm in a thermomixer. Hybridized embryos were then washed in 2 X Stellaris Wash Buffer A for 30 min at 37°. 5 *µ*g/ml of DAPI was added to the final wash for DNA staining. This was followed by 5 min wash with Wash Buffer B (Biosearch Technologies, SMF-WB1-20) and a brief incubation in N-propyl gallate antifade before slide preparation. For slide preparation, an equal volume of N-propyl gallate antifade containing resuspended hybridized embryos and Vectashield antifade were mixed onto the slide. smFISH image stacks were acquired using a DeltaVision Elite inverted microscope on a Photometrics Cool Snap HQ2 camera, with an Olympus PLAN APO 60X (1.42 NA, PLAPON60XOSC2) objective, an Insight SSI 7-Color Solid State Light Engine, and SoftWorx software (Applied Precision) using 0.2 *µ*m z-stacks. Deconvolution was done using Deltavision (SoftWorx) software for all images.

### smiFISH in *C. elegans* embryos

Single molecule inexpensive fluorescence in situ hybridization (smiFISH) was performed according to the protocol described in (Parker *et al*. 2021). Using Oligostan (Tsanov *et al*. 2016) or Stellaris Probe Designer https://oligos.biosearchtech.com/Account/Login?return=/stellaris-designer, primary probes were designed and ordered in 25 nmol 96-well format from IDT (Integrated DNA Technologies), diluted in IDTE (pH 8.0) buffer to 100 *µ*M. DNA oligos were custom-designed to be complementary to the transcript of interest. Individual probes were mixed to reach a final concentration of 0.833 *µ*M. Secondary FLAPX probes containing dual fluorophore labeling at 5 *f* ^*′*^ (*x*) and 3 *f* ^*′*^ (*x*) end, either Cal Fluor 610 or Quasar670, from Stellaris LSC. Fresh smiFISH probes were made for every experiment by annealing target-specific DNA oligos to a fluorescently labeled secondary fluorophore. 2 *µ*L of primary probe mixture set with 1 *µ*L of 50 *µ*M FLAPx secondary probe, 1 *µ*L NEB buffer 3 and 6 *µ*L of DEPC-treated water incubated at 85° for 3 min, 65° for 3 min, 25° for 5 min. 2 *µ*L of annealed probe mixtures were used instead of the usual smFISH probe sets as mentioned

### smiFISH in human cell lines

For human cells lines, smiFISH was performed following Stellaris RNA FISH protocol for adherent cells available online at www.biosearchtech.com/stellarisprotocols. In brief, cells cultured on coverslips were fixed with 4% paraformaldehyde (PFA) for 10 min at room temperature (RT), then permeabilized with 70% (vol./vol.) ethanol for a minimum of 1 hr. Cells on coverslips were washed with Wash Buffer A and incubated at room temperature for 2-5 min. Cells were incubated with smiFISH probes suspended (concentrations as mentioned above) in Sterallis Hybridization Buffer for 4-16 hr at 37° in a humidified chamber. Coverslips were washed in Stellaris Wash Buffer A twice at 37° for 30 min, with 5 *µ*g/ml of DAPI added to the second wash to stain DNA. Following a final 5 min wash with Wash Buffer B, cells were incubated in N-propyl gallate antifade solution and prepared for mounting.

### Immunofluorescence protocol on human cells

To visualize proteins, we used immunofluorescence. U2OS cells were cultured on coverslips to nearly 70% confluency. After fixing the cells with 4% paraformaldehyde (PFA) for 10 min at room temperature (RT) and permeabilizing with Triton-X in 1X PBS for 20 min, they were treated with 1:1000 dilution of Anti-Transportin 1/MIP Antibody (Mouse monoclonal, reacts with human TNPO1, Abcam-ab10303) at room temperature for one hour, followed by 5 times 1X PBS washes. Cells were then treated with 1:500 dilution of Alexa Fluor 488 goat anti-mouse antibody (Invitrogen A11029), and incubated at room temperature for 1 hr, followed by 5 times 1X PBS wash for 5 min each at RT. Finally, the cells were washed with 1X PBS containing 5 *µ*g/ml of DAPI for 5 min and incubated in N-propyl gallate antifade before slide preparation.

### smFISH intensity scanning analysis

To quantify smFISH signal intensity across the nuclear membrane, smFISH microscopy images were processed in FIJI and R. Detailed scripts, output, and reports of this analysis are made available at https://github.com/erinosb/basu_imb2_tnpo1. In FIJI, images were 1) deconvolved, 2) sum-projected (7 z-stacks total), 3) selected for a rectangular region of interest centered over nuclear membrane, 4) quantified by signal intensity integrated over the height of the rectangular ROI with respect to the coordinates of the width of the rectangle, and 5) reported. FIJI-generated reports were then imported into R where they were 1) merged, 2) centered on the middle x-coordinate of the rectangular ROI, 3) normalized by individual-embryo-means, and 3) averaged to create lineplots representing the smFISH intensity means with respect to the x-coordinates of the bounding box and shaded ribbons to represent standard deviation. In addition, we calculated a membrane enrichment metric by comparing the integrated, normalized intensity at the middle of the rectangle (x = 0) with the mean integrated, normalized intensity at x = -50 and +50. We log2-transformed this ratio for easier interpretation. We compared membrane enrichment metrics using Wilcoxon Rank-Sum statistics and BH p-value adjustment for multiple-test correction, where appropriate.

## Results and discussion

### *TNPO1* and *TNPO2* mRNA are homogenously distributed in the cytoplasm

The closest human orthologs of *imb-2* are two duplicated paralogs, *TNPO1* and *TNPO2. C. elegans* IMB-2 protein shares more than 50% identity at the amino acid level with both (Fig. 1a, Figure S1), and they all share analogous domains and motifs (Fig. 1b). In *C. elegans*, the *imb-2* mRNA localizes to the periphery of the nucleus in embryonic cells, and its encoded protein concentrates at both the nuclear periphery and interior (Fig. 1c, d, g). These patterns are most dramatic in the early stages, such as the 2-cell and 4-cell stages, when the cells are largest, but they also persist in later embryonic stages as well (Parker *et al*. 2020). To test whether the human orthologs of IMB-2 are also produced from nuclear-periphery localized mRNA transcripts, we used single molecule inexpensive Fluorescence in Situ Hybridization (smiFISH), a low-cost version of smFISH in which multiple fluorescently labeled probes hybridize to a target mRNA of interest (Raj *et al*. 2010; Tsanov *et al*. 2016; Parker *et al*. 2021). We designed smiFISH probesets to hybridize to human *TNPO1* and *TNPO2* mRNA and tested their localization in U2OS cells. *TNPO1* mRNA and *TNPO2* mRNA showed widespread localization throughout the cytoplasm, with no marked accumulation at the nuclear periphery, but with exclusion from the nucleoplasm (Fig 1e, f). Indeed, *TNPO1* and *TNPO2* mRNA exhibited similar localization to GLYCERALDEHYDE 3-PHOSPHATE DEHYDROGENASE (GAPDH), a control mRNA imaged simultaneously using dual-color probes (Fig. 1e). This illustrates that the mRNA localization pattern of *C. elegans imb-2* mRNA was not conserved in its human orthologs.

### The *C. elegans* protein IMB-2 and its human ortholog TNPO1 share conserved localization in and around the nucleus

Given the lack of conservation at the mRNA localization level, we wanted to test the localization of human TNPO1 protein. To test this, we imaged the TNPO1 protein using immunofluorescence with an anti-TNPO1 antibody. Interestingly, we observed that the TNPO1 protein accumulated in the nucleoplasm and in an area surrounding the nucleus, but it did not tightly co-localize with *TNPO1* mRNA (Fig. 1h). It is striking that the human TNPO1 protein and *C. elegans* IMB-2 protein had similar localization (Fig. 1g, h) despite arising from differing mRNA patterns (*C. elegans imb-2* mRNA at the nuclear envelope and *TNPO1* mRNA in the cytoplasm) (Fig. 1c-f). This suggests that *C. elegans* IMB-2 and human TNPO1 may have different translation dynamics but that the ultimate protein localization was conserved.

### *TNPO1* mRNA is homogenously distributed in the cytoplasm of both U2OS and HeLa cells

To determine whether the homogeneous localization of *TNPO1* mRNA was unique to U2OS cells, we performed *TNPO1* mRNA imaging using smFISH in HeLa cells (Fig. 2a). These cells shared the distributed, homogeneous, cytoplasmic localization of *TNPO1* mRNA seen in U2OS cells.

**Figure 2.**
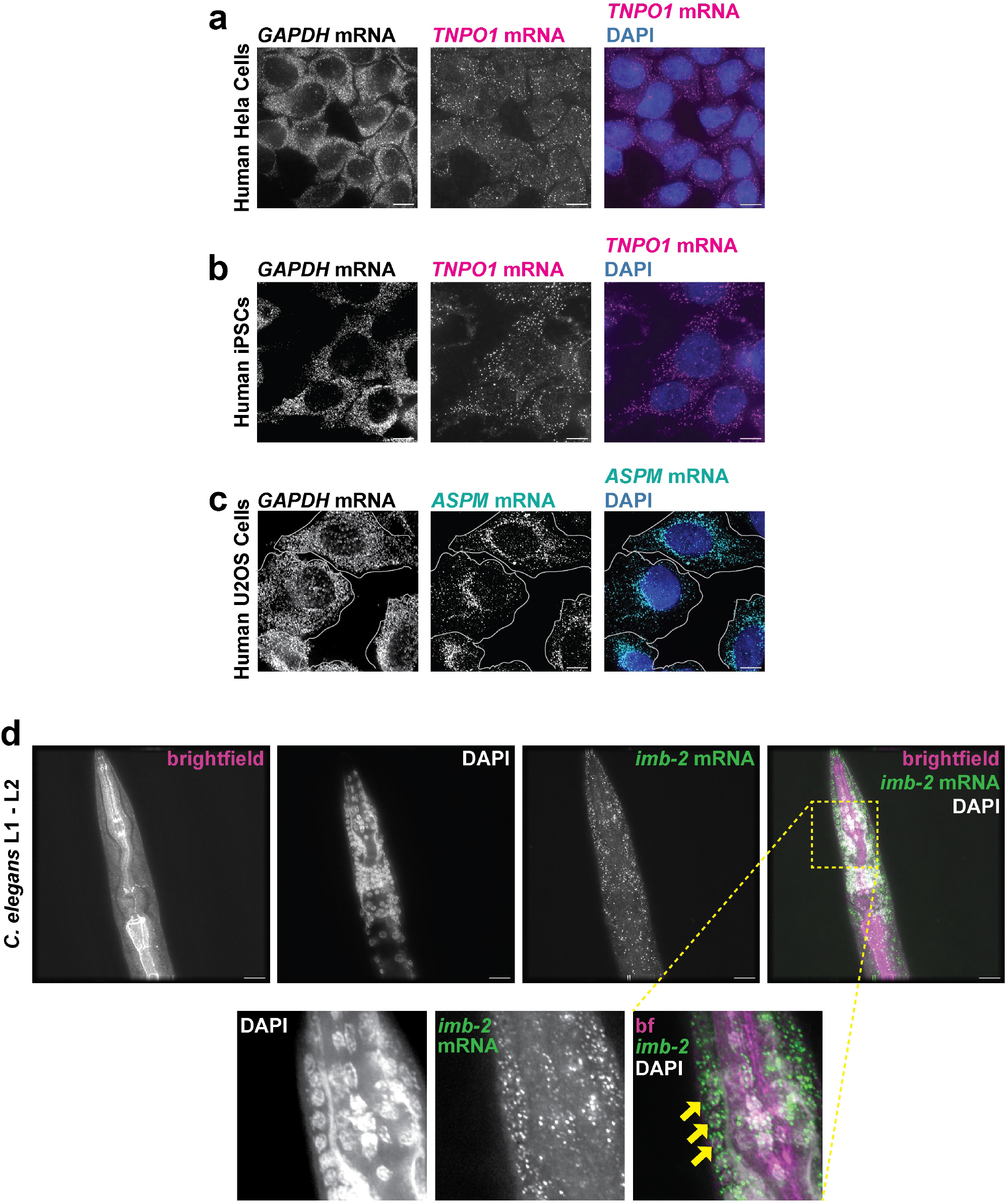
*TNPO1* mRNA patterning in HeLa and induced Pluripotent Stem Cells (iPSCs) phenocopies U2OS expression. **a**. Representative images of human HeLa cells visualizing *TNPO1* (magenta, smiFISH) and nuclei (blue, DAPI) with *GAPDH* (white, smiFISH) included as a homogenous control, Bars 10 *µ*m. **b**. Representative images of human Pluripotent Stem cells (hPSCs) visualizing *TNPO1* (magenta, smiFISH) and nuclei (blue, DAPI) with *GAPDH* (white, smiFISH) included as a homogenous control. Bars 10 *µ*m c. Representative images of *ASPM* mRNA (cyan, smiFISH), a known nuclear periphery-localized transcript, and nuclei (blue, DAPI) with *GAPDH* (white, smiFISH) included as a homogenous control. Bars 10 *µ*m d. Representative images showing *C. elegans imb-2* mRNA (magenta, smFISH), and nuclei (blue, DAPI) in L1 – L2 stage larval worms of ML1651 strain in which the adherens junctions are illuminated by DLG-1::RFP. Bars 10 *µ*m.

### *ASPM* mRNA does localize to the nuclear periphery

As a positive control, we tested ABNORMAL SPINDLE-LIKE MICROENCEPHALY-associated (ASPM), a confirmed mRNA that localizes to the “nuclear edge” in interphase cells and to the peri-centriolar material in mitotic cells (Chouaib *et al*. 2020). Indeed, *ASPM* mRNA localized to nuclear edges in interphase U2OS cells (Fig. 2b), confirming its localization to these structures and providing a valuable control and contrast to *TNPO1* and *TNPO2* mRNA.

### The distinct mRNA localization patterns of *C. elegans imb-2* and human *TNPO1* are species-specific not stage-specific

It is possible that the localization of *C. elegans imb-2* mRNA to the nuclear periphery may be a transient, embryonic-specific behavior, irrelevant in the U2OS cell line which is of osteosarcoma origin. To survey a broader range of developmental stages, we performed *TNPO1* mRNA imaging using smiFISH in human induced Pluripotent Stem Cells (iPSCs) that more closely mirror embryonic states (Fig. 2c). However, they also exhibited homogeneous localization patterns seen in U2OS and HeLa cells. In addition, we expanded *C. elegans imb-2* smFISH to image later stages of the L1 – L2 larval stages. Larvae also showed *imb-2* mRNA localization at the nuclear periphery, confirming that this localization pattern was species-specific, not stage-specific (Fig. 2d). Notably, the patterning was the most visually striking and distinct in early embryos owing to their larger sizes and clearer boundaries.

### The protein localization if IMB-2 and TNPO1 are conserved despite their differing mRNA localization patterns

Overall, we have shown that the mRNA encoding *C. elegans* transportin IMB-2 protein concentrates at the nuclear envelope, suggesting its local translation at this surface. In contrast, the human *TNPO1* mRNA did not accumulate at the nuclear envelope. Still, *C. elegans* IMB-2 protein and its human orthologs all ultimately achieved a similar patterning in the cell – accumulating within and around the nucleus. This emergent conservation is reminiscent of the observation that related yeasts can buffer mRNA abundance differences with compensatory translational efficiency differences, thereby preserving conservation at the proteome level (McManus *et al*. 2014). In the case of human Tnpo1, it is likely that post-translational protein transport plays a larger role in its localization than local translation.

Interestingly, the nuclear envelope is known to be a site of local translation, as a subset of yeast nucleoporins and nuclear pore complex components colocalize with their own mRNAs (Lautier *et al*. 2021; Penzo and Palancade 2023). Therefore, though the exact orthologs that undergo local translation may not be conserved, certain subcellular locales may potentially act as “hot spots” for local translation. This may be required to ensure their proteins fold properly, coordinate with the membrane, or associate with membrane complexes in the proper order or at the proper interface.

## Data availability

Strains and plasmids are available upon request. Figure S1 contains all raw data, processed data, and statistical analyses used in *C. elegans* microscopy quantifications. Supplemental Table 1 contains all raw data, processed data, and statistical analyses used in human cell microscopy quantifications. Supplemental Table 2 contains all probeset sequences used in this study. Code used to quantify and plot all figures is available at https://github.com/erinosb/basu_imb2_tnpo1.

## Acknowledgments

This work utilized resources from the University of Colorado Boulder Research Computing Group, which is supported by the National Science Foundation (awards ACI-1532235 and ACI-1532236), the University of Colorado Boulder, and Colorado State University. This work utilized microscopy resources from the CSU Microscope Imaging Network. Some strains were provided by the Caenorhabditis Genetics Center, which is funded by the National Institutes of Health Office of Research Infrastructure Programs (P40 OD010440). Some figure elements were created in BioRender. We are grateful to Tim Stasevich, Gabriel Galindo, and O’Neil Wiggan for their assistance with U2OS and HeLa cell culture protocols. We are also grateful to Dr Kevin Flynn and Jillian Kuizenga to provide us with iPSCs.

## Supplemental Figures

**Table 1.**
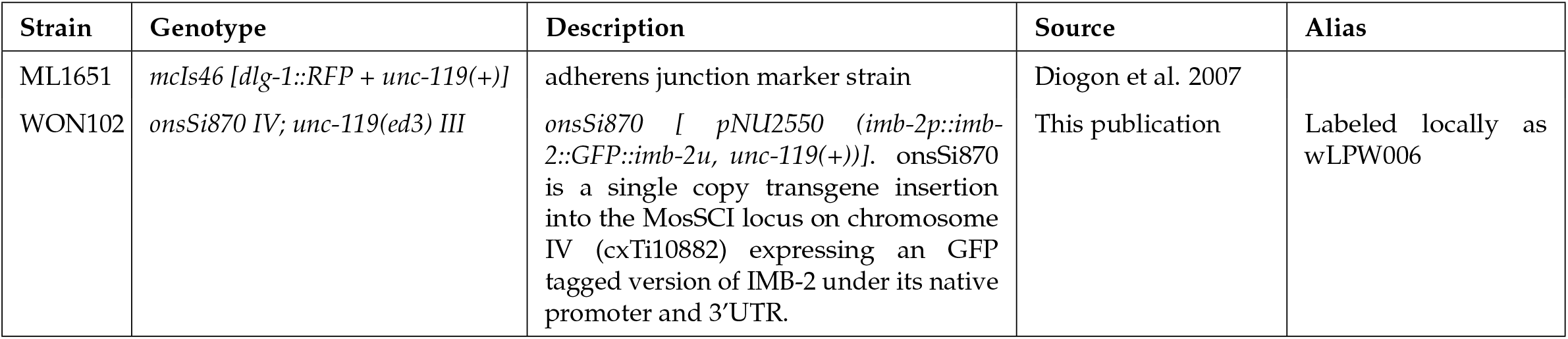
Strains used in this study.

**Figure S1.**
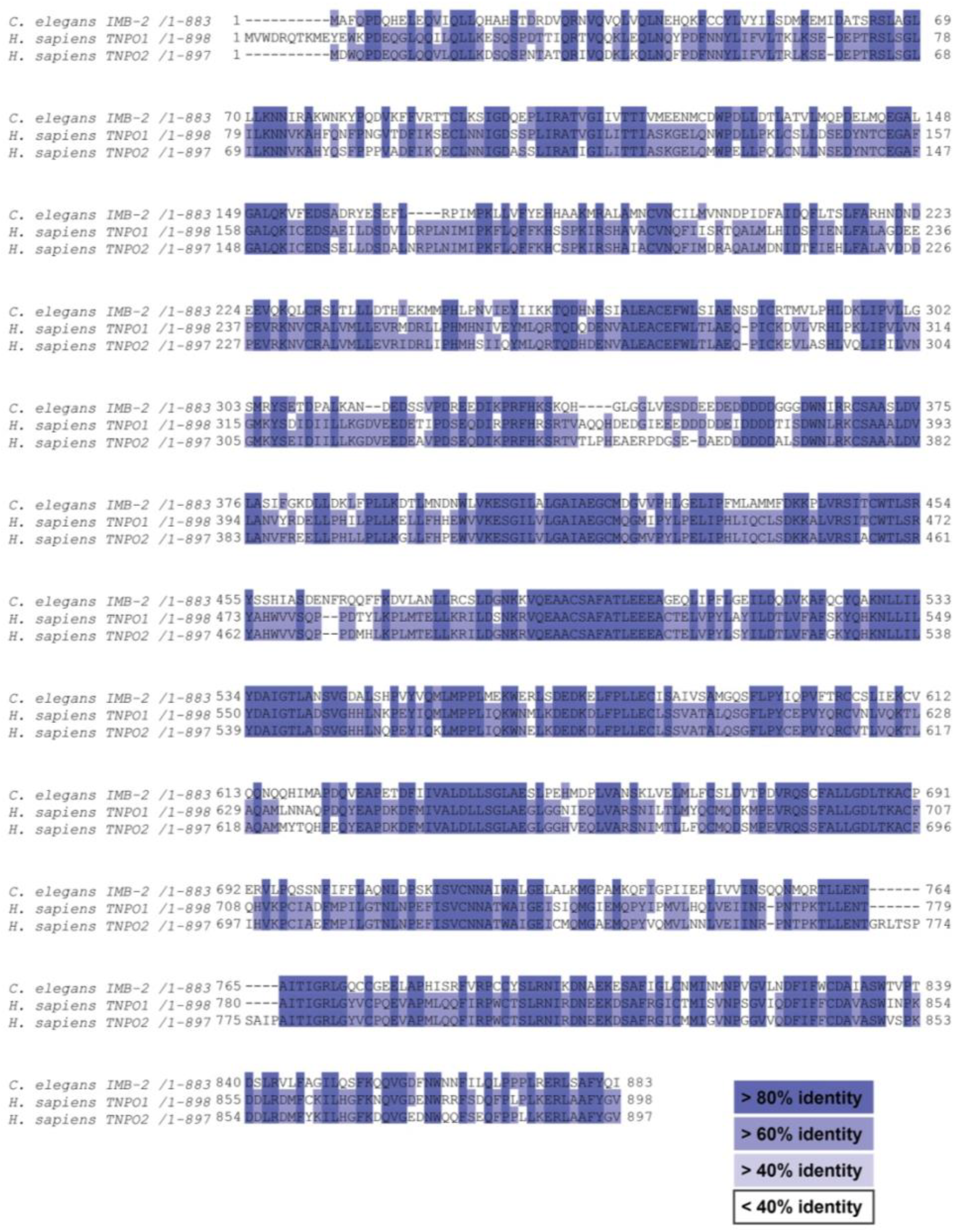
Multiple sequence alignment of *C. elegans* IMB-2, human Tnpo1, and human Tnpo2 proteins. Alignments as generated by Clustal Omega. Colors represent percent identity at the amino acid level.

